# Transcriptional profiling from whole embryos to single neuroblast lineages in *Drosophila*

**DOI:** 10.1101/2022.04.04.487012

**Authors:** Austin Seroka, Sen-Lin Lai, Chris Q Doe

## Abstract

Embryonic development results in the production of distinct tissue types, and different cell types within each tissue. A major goal of developmental biology is to uncover the “parts list” of cell types that comprise each organ. Here we perform single cell RNA sequencing (scRNA-seq) of the *Drosophila* embryo to identify the genes that characterize different cell and tissue types during development. We assay three different timepoints, revealing a coordinated change in gene expression within each tissue. Interestingly, we find that the *elav* and *mhc* genes, whose protein products are widely used as markers for neurons and muscles, respectively, show broad pan-embryonic expression, indicating the importance of post-transcriptional regulation. We next focus on the central nervous system (CNS), where we identify genes characterizing each stage of neuronal differentiation: from neural progenitors, called neuroblasts, to their immediate progeny ganglion mother cells (GMCs), followed by new-born neurons, young neurons, and the most mature neurons. Finally, we ask whether the clonal progeny of a single neuroblast (NB7-1) share a similar transcriptional identity. Surprisingly, we find that clonal identity does not lead to transcriptional clustering, showing that neurons within a lineage are diverse, and that neurons with a similar transcriptional profile (e.g. motor neurons, glia) are distributed among multiple neuroblast lineages. Although each lineage consists of diverse progeny, we were able to identify a previously uncharacterized gene, *Fer3*, as an excellent marker for the NB7-1 lineage. Within the NB7-1 lineage, transcriptional clusters are identifiable in neuroblasts and neurons, and each cluster is composed of current temporal transcription factor (e.g. Hunchback, Kruppel, Pdm, and Castor), novel temporal factors, and/or targets of the temporal transcription factors. In conclusion, we have characterized the embryonic transcriptome for all major tissue types and for three stages of development, as well as the first transcriptomic analysis of a single, identified neuroblast lineage, finding a lineage-enriched transcription factor.

## Background

Understanding how tissues such as the nervous system develop is a central goal of developmental biology. An important part of development is the generation of cell types that vary in molecular profile, cell morphology and cell function. Identifying the different cell types in a tissue is particularly important for the study of neural development, where a vast number of distinct neurons must interconnect to form a functional nervous system (Luo, 2020). Defining neural diversity at the molecular level is important for generating a “parts list” of neurons in the brain, and will ultimately generate a better understanding of how neuronal diversity is established. Increased understanding of the transcriptional mechanisms used to generate the appropriate neurons in the correct spatial location and correct time also advances the potential for neurotherapeutics to counteract injury or disease.

Single cell RNA-sequencing (scRNA-seq) is a powerful method for determining the transcriptional profile of complex tissues such as the central nervous system (CNS), including mammalian cortical excitatory and inhibitory neurons (Shen et al., 2020), hippocampal neurons (Hodge et al., 2019), zebrafish embryonic neurons (Tambalo et al., 2020), and Drosophila larval, pupal and adult neurons (Brunet Avalos and Sprecher, 2021; Davie et al., 2018; Konstantinides et al., 2018; McLaughlin et al., 2021; Nguyen et al., 2021; Vicidomini et al., 2021; Xie et al., 2021) (Velten et al., 2022). Surprisingly, there has yet to be a sc-RNAseq study of Drosophila embryonic neurogenesis; the only embryonic sc-RNAseq study was tightly focused study on pre-gastrula embryos (Karaiskos et al., 2017).

*Drosophila* neurogenesis is ideal for the application of transcriptional analysis, as there is a wealth of cell- and tissue-specific genes that provide ground-truth information for identifying cell types through the use of sc-RNAseq. In the *Drosophila* embryo, neuronal diversity is generated in three steps: (1) spatial patterning cues are used to generate neural progenitor (neuroblast) identity, with each neuroblast having a unique identity based on its spatial location (Skeath and Thor, 2003); (2) temporal patterning generated by a cascade of “temporal transcription factors” (TTFs; Hunchback > Kruppel > Pdm > Castor) diversifies ganglion mother cell (GMC) identity within each neuroblast lineage (Doe, 2017); and (3) nearly all GMCs undergo a terminal asymmetric division to partition the Notch inhibitor Numb into one of the two siblings, thereby creating Notch^ON^ and Notch^OFF^ siblings that have unique molecular and morphological identities (Mark et al., 2021; Skeath and Doe, 1998; Truman et al., 2010).

Here we present a scRNA-seq atlas of the entire embryo at three timepoints. We subsequently focus on gene expression changes within the developing nervous system: first at stage 12 when neuroblasts are maximally proliferative and only the earliest-born neurons have begun to differentiate; then at stage 14 when both neuroblasts and differentiated neurons are well represented; and finally at stage 16 where the bulk of the mature embryonic neurons are present. In addition to tracking the transcriptome of bulk neuroblasts, GMCs and neurons, we also address the question of whether sc-RNAseq can be used to identity lineage-specific gene expression. Here, we focus on the best characterized neuroblast lineage: NB7-1 which is a model for studying spatial patterning (McDonald et al., 1998), temporal patterning (Isshiki et al., 2001; Kohwi et al., 2013; Meng et al., 2019; Meng and Heckscher, 2020; Pearson and Doe, 2003; Seroka et al., 2020; Seroka and Doe, 2019), and Notch^ON^/Notch^OFF^ sibling specification (Mark et al., 2021; Skeath and Doe, 1998). We can identify two classes of motor neurons and interneurons known to be present in this lineage, as well as novel gene expression patterns that identify candidates for lineage-specific functions. To our knowledge, our study is the first to characterize the Drosophila post-blastoderm embryonic transcriptome and the first to transcriptional profile a single neuroblast lineage.

## Results and Discussion

### The transcriptome of all embryonic cell types

To create a transcriptional time course of embryonic development, we dissociated stage 12, 14, and 16 embryos using standard methods and independently performed two technical replicates for 10X Genomics scRNAseq on cells from each timepoint. From stage 12 embryos we isolated 20,038 cells and obtained at 1,234 median genes per cell; from stage 14 embryos we isolated 28,045 cells at 656 median genes per cell; and from stage 16 embryos we isolated 24,032 cells at 450 median genes per cell. Cells were filtered for quality in Seurat using standard methods. Following quality control, the stage 12, 14 and 16 objects contained 17,564, 24,668 and 20,328 cells respectively. We merged these datasets using Seurat to obtain a single UMAP atlas containing 62,560 cells and 96 clusters (Figure 1A; Supplemental Table 1).

**Figure 1.**
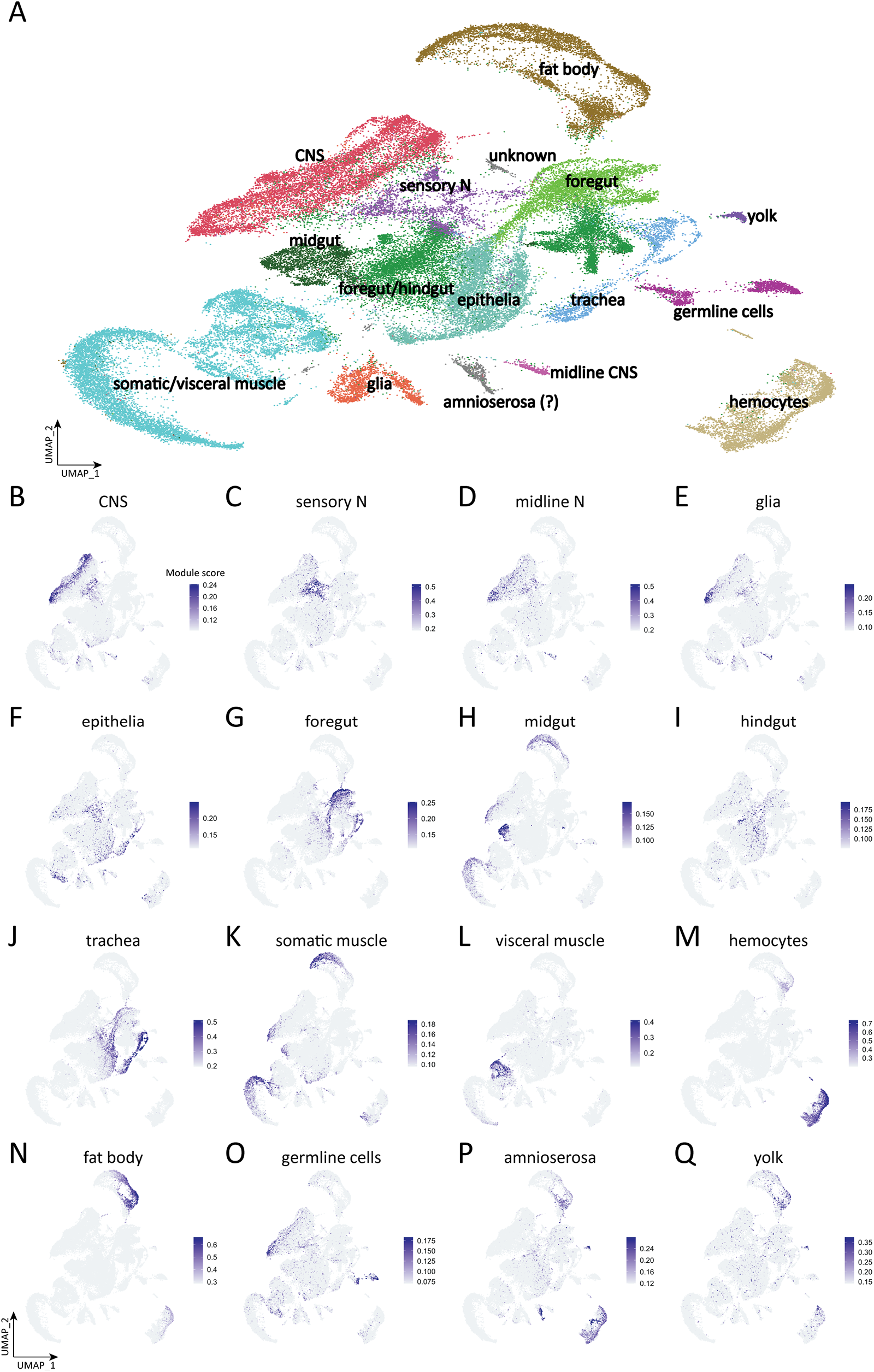
Tissue type atlas of whole *Drosophila* embryo. (A) Integrated single-cell atlas of whole *Drosophila* embryos. Cluster identity is determined by the module scores based on the tissue-specific genes. (B-Q) Plots of module scores of tissue-specific genes for each individual tissue in UMAP space. The colors encode the module scores computed by AddModuleScore function in Seurat. Tissue defining genes listed in Supplemental Table 2.

We observed clusters representing all expected embryonic cell types, identified by tissue-specific annotations in Flybase (www.flybse.org) and BDGP in situ atlas (https://insitu.fruitfly.org/cgi-bin/ex/insitu.pl) (Tomancak et al., 2007). For example, genes annotated as “glia” were used to score cells based on their expression as shown in Figure 1E, and the scores were used to assign the “glial” distribution in the UMAP object; a similar process was done for all tissue types. The UMAP for each annotated pool of genes is shown in Figure 1B-Q, and the list of genes in each pool is given in Supplemental Table 2.

Clusters included neural cell types in the central nervous system (Figure 1B), ciliated sensory neurons (Figure 1C), midline cells (Figure 1D), and glia (Figure 1E). These neural cell types will be further subdivided and characterized in more depth below.

We also observed clusters representing epithelia (Figure 1F); foregut (Figure 1G); midgut (Figure 1H); hindgut (Figure 1I); trachea (Figure 1J); somatic/visceral muscle (Figure 1 K, 1L); hemocytes (Figure 1M); fatbody (Figure 1N); germline cells (Figure 1O); amnioserosa (Figure 1P); and yolk (Figure 1Q). In addition to identifying all expected embryonic cell types, we also observe clusters that do not express tissue-specific genes (unknown, Figure 1A); these could be previously uncharacterized cell types or a mixture of cell types that were not well clustered. We conclude that we have identified transcriptional profiles for all major embryonic cell types.

### Tissue-specific proteins can be widely transcribed

We were surprised to see widespread non-neural expression of the *embryonic lethal abnormal vision (elav)* gene, whose protein product Elav is widely used as a marker for post-mitotic neurons in Drosophila (Robinow and White, 1991) and vertebrates (Park et al., 2000). It has previously been noted that *elav* is transcribed in proliferating neuroblasts but not translated (Berger et al., 2007); we confirm here strong *elav* transcription in the neuroblast clusters (Figure 2A). Surprisingly, we also found *elav* broadly transcribed at lower levels in all tissue types of the embryo, including mesodermal derivatives, glia, trachea,gut, and fat body (Figure 2A). An *elav* paralog, *found in neurons (fne)*, also shows the same pattern of high-level expression in neuronal clusters and broad, lower-level expression in all cell types (Figure 2B). This suggests that only post-mitotic neurons have a mechanism for translating the *elav* and *fne* transcripts. Similarly, the C. elegans single ortholog of Elav, named Exc-7,is also expressed in non-neuronal cell types including muscle and hypoderm (Pham and Hobert, 2019). In contrast, none of the zebrafish orthologs show noticeable transcription outside the CNS (Farnsworth et al., 2020).

**Figure 2.**
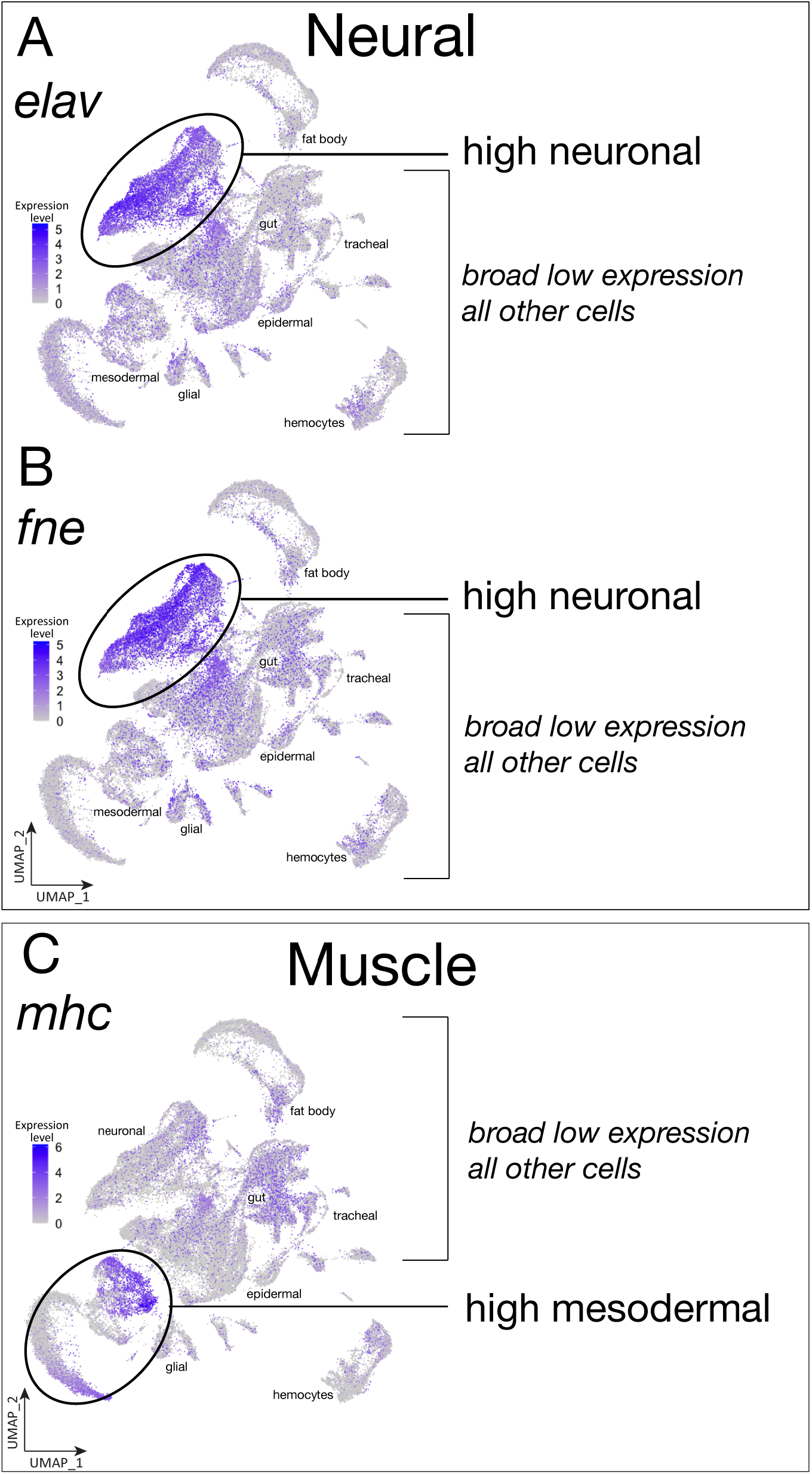
Tissue-specific proteins can be widely transcribed. (A) Plot of *elav* expression level in whole embryo single-cell atlas. (B) Plot of *fne* expression level in whole embryo single-cell atlas. (C) Plot of *mhc* expression level in whole embryo single-cell atlas.

To see if this mechanism could be generalized to another tissue, we examined several pan-mesodermal genes, and while most showed narrow expression in some or all mesodermal derivatives (Figure 1C), the *myosin heavy chain (mhc)* gene showed broad expression in all cell types (Figure 2C), even though Mhc protein is only detected in mesodermal lineages. We conclude that some genes are transcribed widely followed by tissue-specific translation. Candidates for positive regulators of this process would be RNA-binding proteins or long non-coding RNAs enriched specifically in mesodermal derivatives (Supplemental Table 1). It will be of interest to understand the global mechanisms regulating mRNA translation that refine these broad expression pattern to unique cell types. We conclude that some tissue-specific or cell type-specific proteins are widely transcribed, revealing a major role for post-transcriptional regulation. It will be interesting to determine the mechanism of the post-transcriptional regulation either via RNA-binding proteins or microRNAs.

### Developmental timeline of all embryonic cell types

In order to visualize the developmental trajectories of each identified cell type in the atlas, we plotted the timepoint of origin of each cell (Figure 3). We observe that unbiased clustering orders most cells of each identity along a maturation axis from stage 12 to stage 16. In some cases (CNS neuroblasts, germline cells,yolk) we observe less transcriptional differences over time, with cells from each timepoint clustering together instead of separating along a developmental axis. We draw three conclusions from these data: Firstly, most tissue types are established early in embryogenesis and change their transcriptional programs as they mature over time. Secondly, cell types such as CNS and glial progenitors continually produce progeny and show less variability along a developmental axis, as their core transcriptional identity as progenitors is maintained from stages 12-16. Thirdly, cells in the same tissues may develop synchronously, like fat body or hemocytes, so the cells at different developmental stages cluster together to form developmental trajectories. In contrast, the cells that develop asynchronously such CNS or epithelia do not cluster together during development. Lastly, some cell types such as germline and yolk cells are established early in development and their transcriptomes stay constant across time with almost no change (Figure 3).

**Figure 3.**
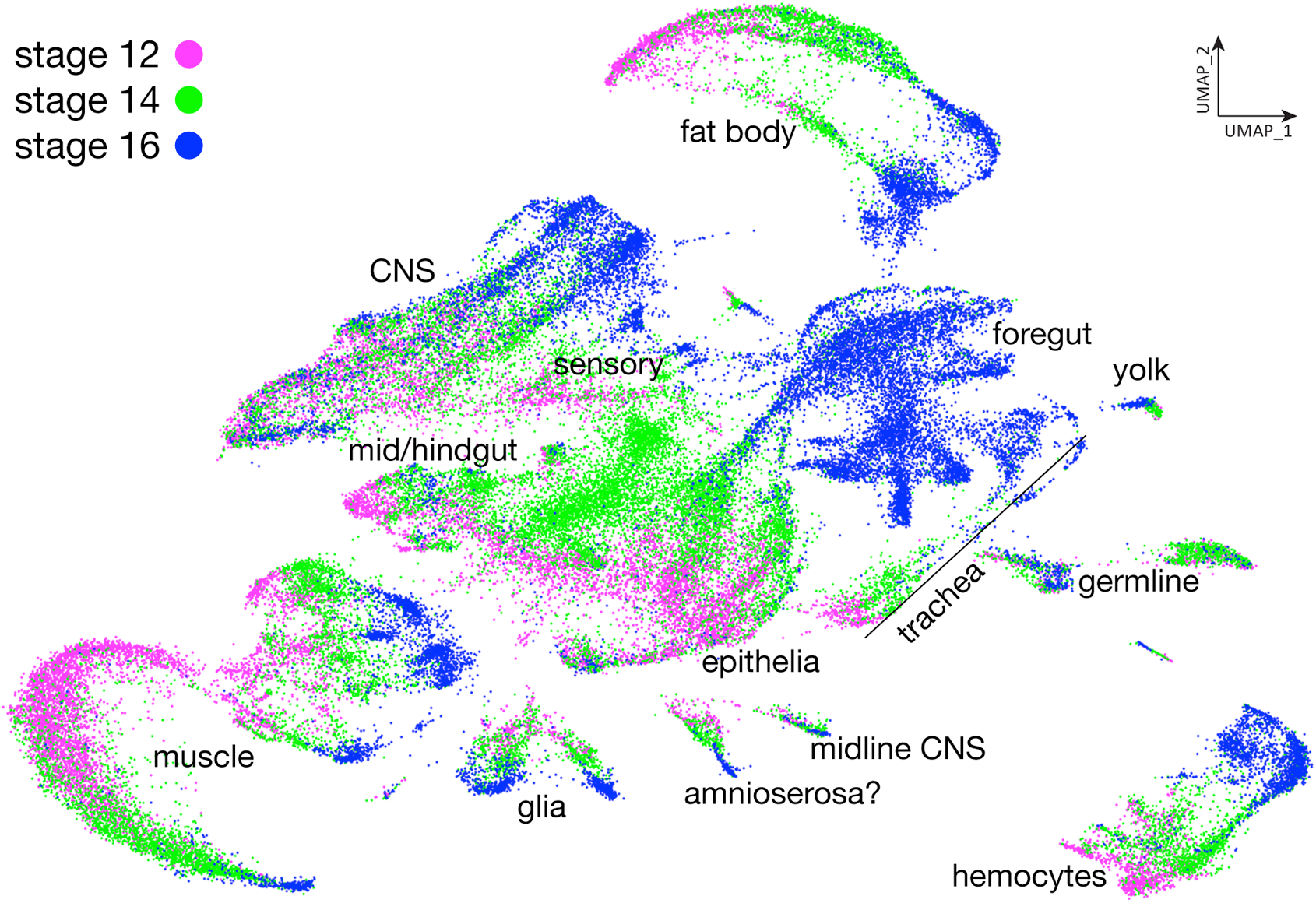
Developmental trajectory of all embryonic cells. Plot of integrated single cell atlas based on their developmental stages in UMAP space. Stage 12, magenta; stage 14, green; stage 16, blue.

### Neural cell type atlas

We next wanted to characterize the neural transcriptomes in more detail, so we extracted the neuronal clusters (4, 13, 14, 22, 23, 25, 27, 28, 37, 39, 45, 47, 53, 59, 61, 67, 71,73, 75, 85) from the merged all cells data set shown in Figure 1A and generated a new “embryonic neural cell type” atlas containing 13,136 cells distributed into 33 clusters (Supplemental Table 3). We then manually assigned different CNS cell types (Figure 4) based on the following experimentally validated marker genes: neuroblasts, *miranda* (*mira*) (Figure 4B); GMCs, *tap* (Figure 4C); new-born Notch^ON^ neurons, *Hey* (Figure 4D); young neurons, *neuronal synaptobrevin* (*nSyb*)+ *bruchpilot* (*brp*)-(Figure 4E); mature neurons (or old neurons), *brp* (Figure 4F); midline cells, *single-minded* (*sim*) (Figure 4G); sensory neurons, *Root* (Figure 4H); glia, *reverse potential*(*repo*) (Figure 4I); and possibly enteric neurons, *Ecdysone-inducible gene E2* (*ImpE2*) (Figure 4J). We conclude that our merged stage 12, 14, and 16 atlas has representation from all major neural cell types.

**Figure 4.**
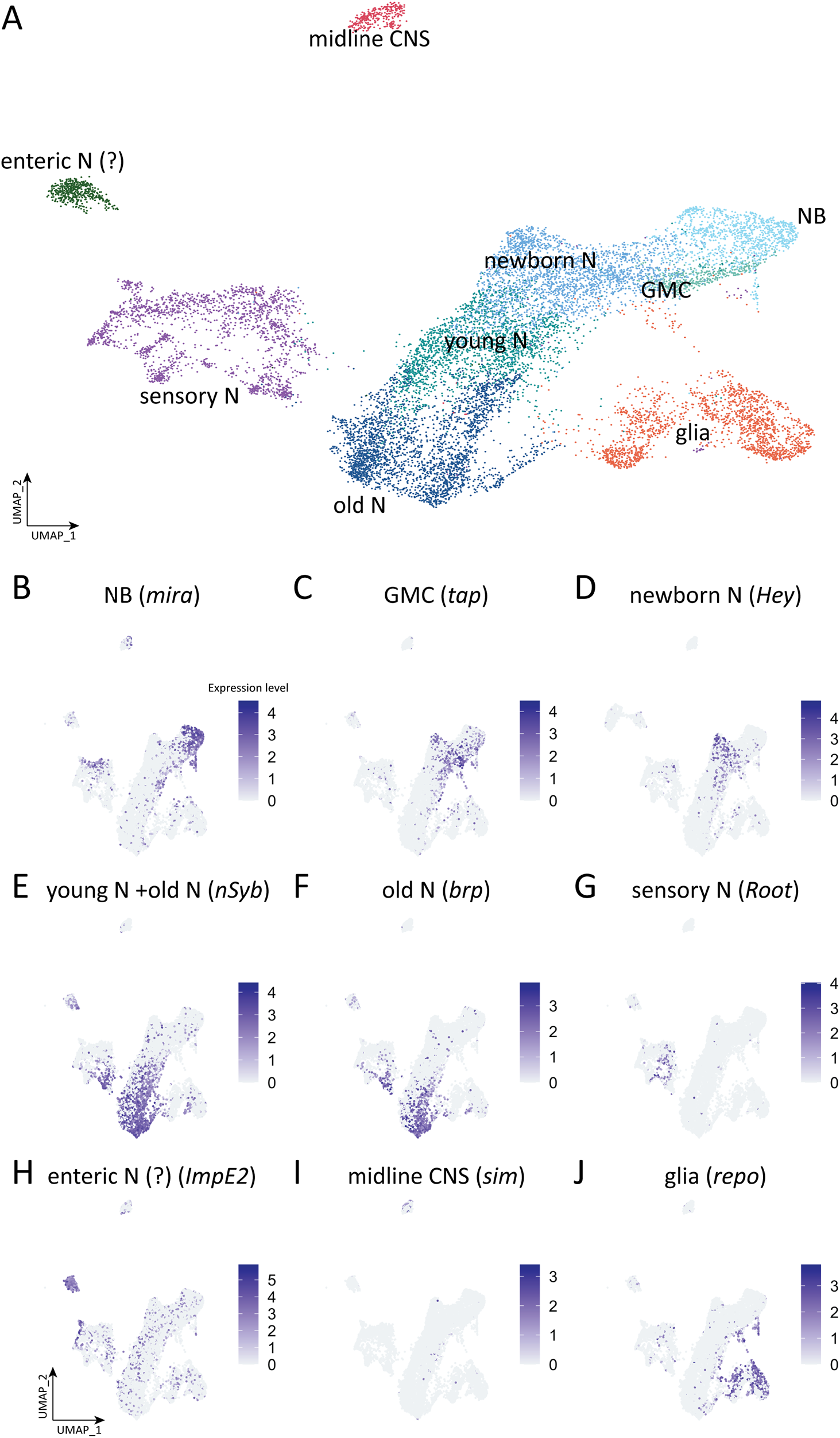
Atlas of reclustered embryonic nervous system. (A) Integrated single-cell atlas of *Drosophila* embryonic nervous system. Cluster identity is determined by the expression ground-truth genes as indicated. (B-J) Plots of expression level of ground-truth genes for each individual neural cell type in UMAP space. Colors encode logarithm-transformed expression levels.

We next narrowed our focus to the CNS only, and asked how the cell type-specific clusters changed over the three developmental stages analyzed here (stages 12, 14, 16). Stage 12 embryos are actively undergoing neuroblast divisions in this early stage of neurogenesis, while by stage 14 neurogenesis and axon outgrowth are proceeding. By stage 16 neurons are actively involved in axon guidance, dendrite outgrowth, and synaptic connectivity (Goodman and Doe, 1993). Confirming previous work, we find that there is a general shift from expression of neuroblast markers to mature neuronal markers across these timepoints (Figure 5A; Table 1). For example, the neuroblast marker *miranda* (*mira*) is expressed in many cells at stage 12, but few at stage 16 (Figure 5B; Table 1); conversely, the mature neuron marker *brp* is barely detected at stage 12 (these may be pioneering motor neurons (Thomas et al., 1984)), but broadly expressed at stage 16 (Figure 5F; Table 1). Genes characterizing other stages of neuronal development fall in between these extremes (Figure 5C-E; Table 1). We conclude that cell type specific clusters validated by ground truth experimental data for cluster defining genes will provide a rich resource for further identification and functional characterization that generate cell type-specific biology (e.g. neuroblast self-renewal or asymmetric division in the neuroblast cluster, or synaptogenic molecules in the mature neuron cluster). For cell types with few markers, such as GMCs or young neurons, our atlas provides the opportunity to identify additional cell type-defining genes.

**Figure 5.**
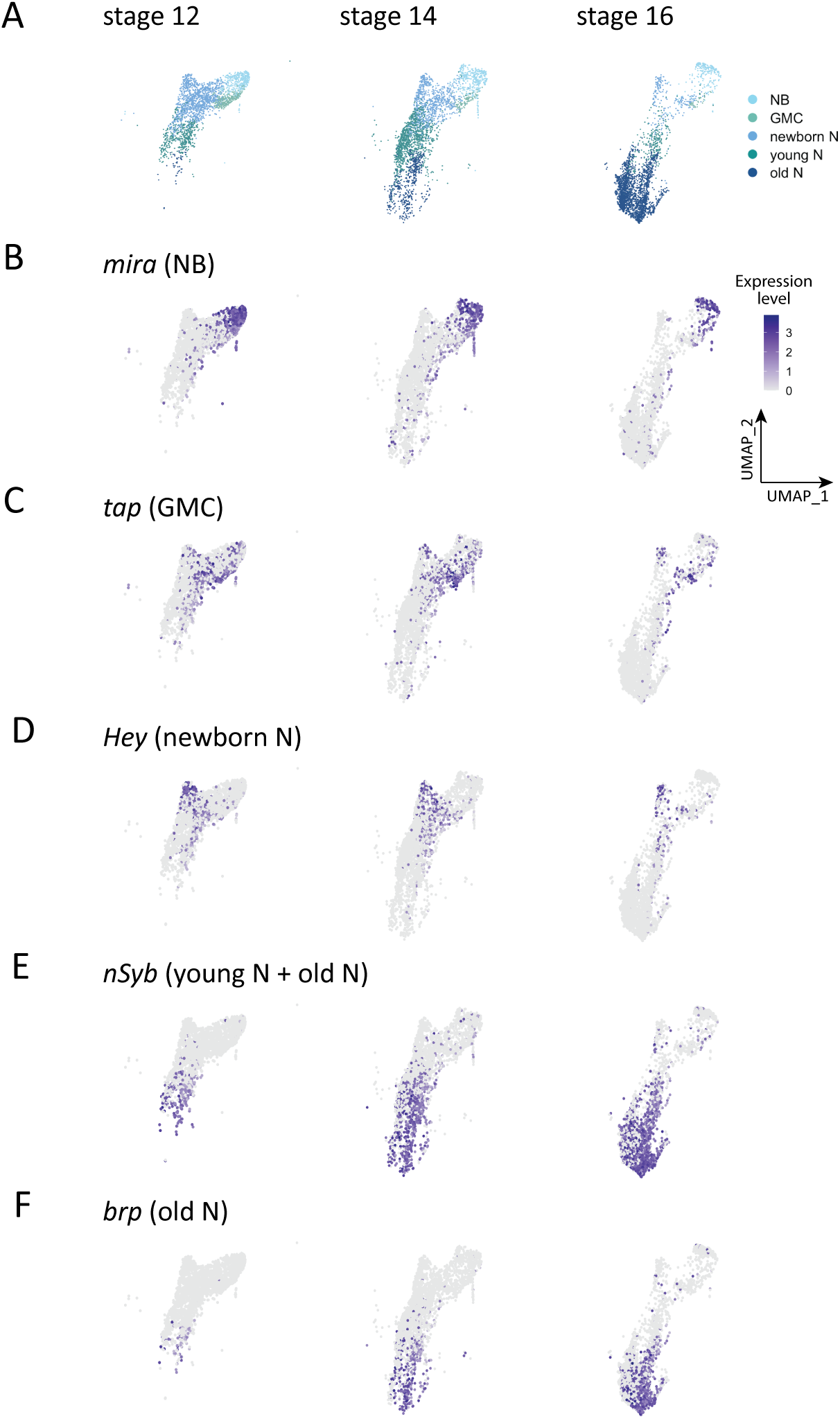
Neural cell type atlas. (A) Atlas of neural cells at embryonic stage 12, stage 14, and stage 16. NB, neuroblast; GMC, ganglion mother cell; newborn N, newborn neuron; young N, young neuron; old N, mature neuron. (B-F) Distribution of ground-truth genes in the neural cell UMAP at embryonic stage 12, stage 14, and stage 16. Colors encode logarithm-transformed expression levels.

**Table 1.** Cell types identified by specific gene expression across development.

|  |  |  |  |
| --- | --- | --- | --- |
|  | Stage 12<br>(2589 cells) | Stage 14<br>(3041 cells) | Stage 16<br>(2490 cells) |
| NB | 18.7% | 12.2% | 8.5% |
| GMC | 11.6% | 3.4% | 1.2% |
| Newborn N | 49.8% | 27.3% | 9.0% |
| Young N | 18.0% | 39.6% | 10.2% |
| Old N | 2% | 17.4% | 71.1% |

### Neural cell type clustering of transcription factors and cell surface molecules

The most differentially expressed genes in the larval CNS and in other organisms are transcription factors (TFs) and cell surface molecules (CSMs) (Li et al., 2020). Here we characterize TFs expressed in the neuroblasts at each developmental timepoint (Figure 6A). We first used Seurat to identify the most enriched TFs at each stage, and then calculated the expression levels of TFs combined from stage 12,stage 14, and stage 16. We then used heatmap visualization to show the scaled average expression of each TF at different stages. We found that NBs express different TF subsets over time, but these subsets did not correlate with known temporal TF expression (*hb*, *pdm2*, or *cas*) (Figure 6A). This may be due to lineage asynchrony amongst the total NB population. Of course, it remains possible that different TFs are co-expressed with temporal TFs when examined at a single lineage level of resolution; we explore this possibility in the following section.

**Figure 6.**
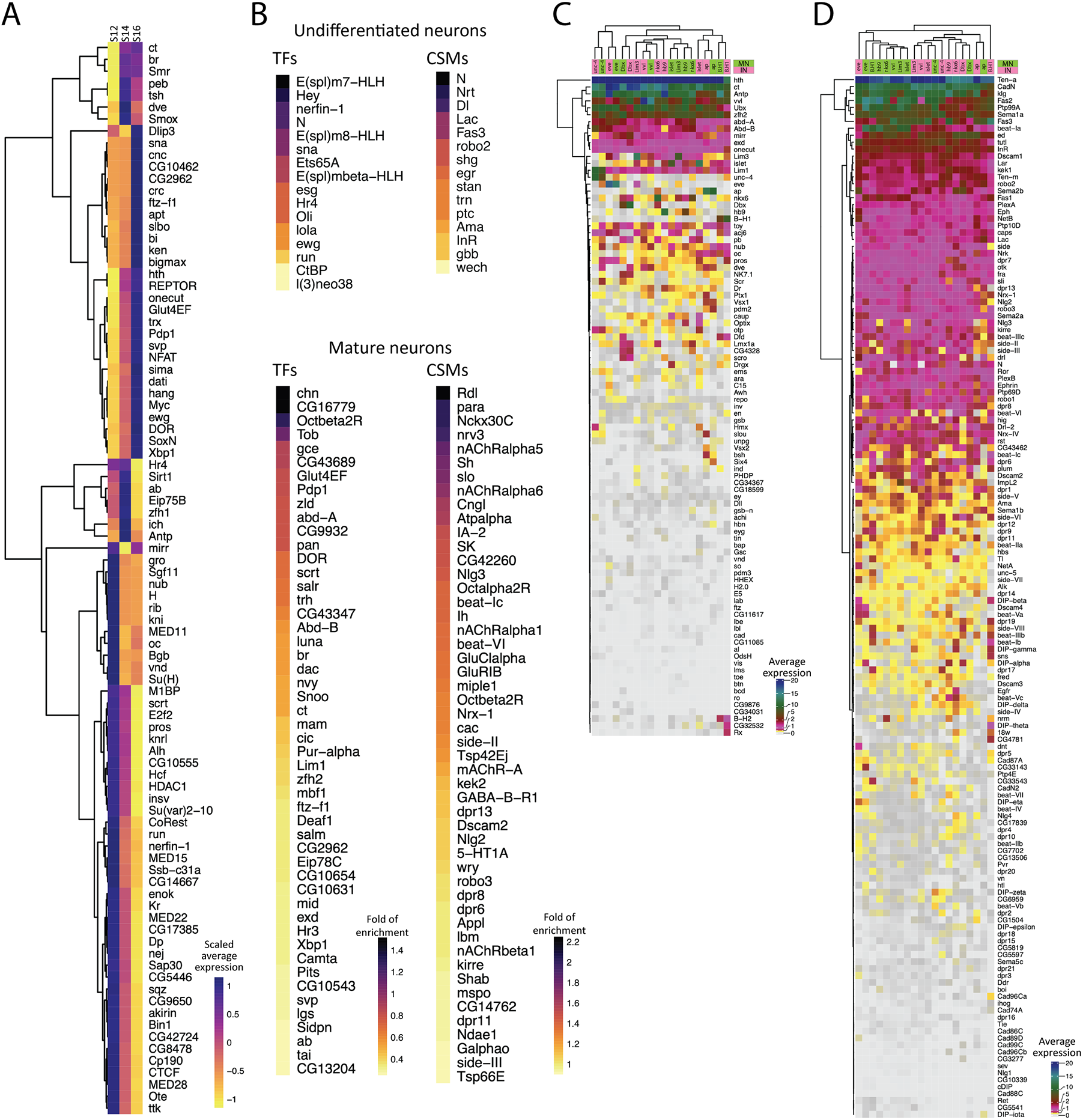
Gene expression profiles in neuroblasts and neurons. (A) Heatmap of scaled average expression of pooled transcription factors statistically enriched in neuroblasts at stage 12 (S12), stage 14 (S14), and stage 16 (S16). Gene names are listed at the right side. Colors encode the levels of scaled average expression of the transcription factors at different stages. Dendrogram at the left shows the clustering of the transcription factors based on the levels of scaled average expression. (B) Heatmap of statistically enriched transcription factors (TFs) and cell surface molecules (CSMs) in undifferentiated and mature neurons. Gene names are listed at the right side. The colors encode the logarithm-transformed folds of enrichment of average expression. (C-D) Heatmap of homeodomain transcription factors (HDTFs) (C) and cell surface molecules (CSMs) (D) in selected motor neurons (MNs) and interneurons (INs). Gene names are listed at the right side. Colors encode the levels of average expression of HDTFs (C) and CMSs (D). Gene names are shown at the right side. Dendrogram at the top shows the clustering of different types of cells, and dendrogram at the left shows the clustering of HDTFs (C) or CSMs (D).

We next switched to analyzing the transcriptomes of undifferentiated cells (*hdc*+) and mature neurons (*brp*+) to understand the differences between immature and fully differentiated neurons (Figure 6B). We focused on TFs and cell surface molecules (CSMs), as these groups of genes are known to be highly differentially expressed in larval neurogenesis (Li et al., 2020). We found undifferentiated cells are enriched for Notch signaling pathway genes (*Dl*, *Hey, E(spl)m7-HLH, E(spl)m8-HLH, E(spl)mβ-HLH*, and *N*), or neuroblast-related genes (*esg, l(3)neo38, run*, and *sna*). In contrast, mature neurons are enriched for TFs promoting cell fate (*ab*, *br*, *ct*, *dac, ftz-f1*, and *zfh2*), and CSMs for synapse formation (*beat IIc, beat-VI, dpr6*, and *dpr8*,) and physiological functions (*nAChRα1*, *nAChRα5, nAChRα6, Octα2R*). We conclude that Notch signaling is important in early neurogenesis, whereas some TFs and CSMs play a greater role in mature neurons. This is consistent with previous findings showing that Notch signaling is important for specifying sibling neurons following GMC division (Skeath and Doe, 1998).

As first shown in *C. elegans*, each neuron expresses a unique code of homeodomain TFs that correlate with, and in some cases specify, their identity (Reilly et al., 2020). Thus, we selected homeodomain TFs (HDTFs) previously shown to be expressed in motor neurons and interneurons, and looked for additional HDTFs that clustered with each of these well-defined neuronal types (Figure 6C). Interestingly, we found that ventral muscle motor neurons (*hb9*+, *islet+, Lim3+*, and *nkx6+)* express similar set of HDTFs, and they are not clustered with dorsal muscle motor neurons (*eve*+) or lateral muscle motor neurons (*B-H1*+). However, *hb9+, islet+, Lim3+*, and *nkx6+* interneurons do not cluster together like the motor neurons. Moreover, motor neurons and interneurons expressing *ap, B-H1*, *Dbx, eve, unc-4*, or *vvl* are clustered together, suggesting that these motor neurons are not very different from the interneurons. Furthermore, all motor neurons and interneurons express more than one HDTF, suggesting that each neuron may expresses a unique set of HDTFs to specify their identity, similar to neurons in C. elegans (Reilly et al., 2020).

We next wanted to identify the CMSs that may be regulated by, and thus co-clustered with, motor neuron expressed HDTFs (Figure 6D). We focused on the CSMs reported in Özkan et al. (Özkan et al., 2013) and the phosphotyrosine kinases/phosphatases. First, we found motor neurons cluster with a similar set of CSMs,regardless of their muscle targets (Figure 6D). Despite this observation, ventral muscle motor neurons (*hb9+, islet+, Lim3+*, and *nkx6+)* remain clustered independently of the dorsal muscle MNs (*eve*+) and lateral muscle MNs (*B-H1*+) (Figure 6D). In addition, interneurons express similar set of CSMs, and cluster together distinctly from motoneurons, with the exception of *eve+* and *B-H1+* interneurons, which cluster away from other interneurons (Figure 6D). Secondly, different motor neurons and interneurons use different combination of CSMs (Figure 6D). It remains unclear if the activation of a CSM requires only one HDTF or a unique combination of HDTFs.

### NB7-1 single lineage gene expression profiles across embryonic development

To our knowledge, transcriptional profiling of an individual neuroblast lineage has not yet been performed. Here we identify the NB7-1 lineage by using NB7-1-gal4 to drive the expression of a RedStinger reporter, which is identifiable *in-silico* by subsetting cells expressing the RedStinger transcript (Figure 7A). We use this lineage-specific transcriptome to address these questions. (1) Do the cells in the NB7-1 lineage cluster together? (2) Can we identify lineage-specific genes, that could maintain the spatial identity of the NB throughout development? (3)Does NB7-1 undergo the same pattern of embryonic stage-specific differentiation? (4) Is it possible to detect genes co-clustered with the temporal TFs within a single NB lineage?

**Figure 7.**
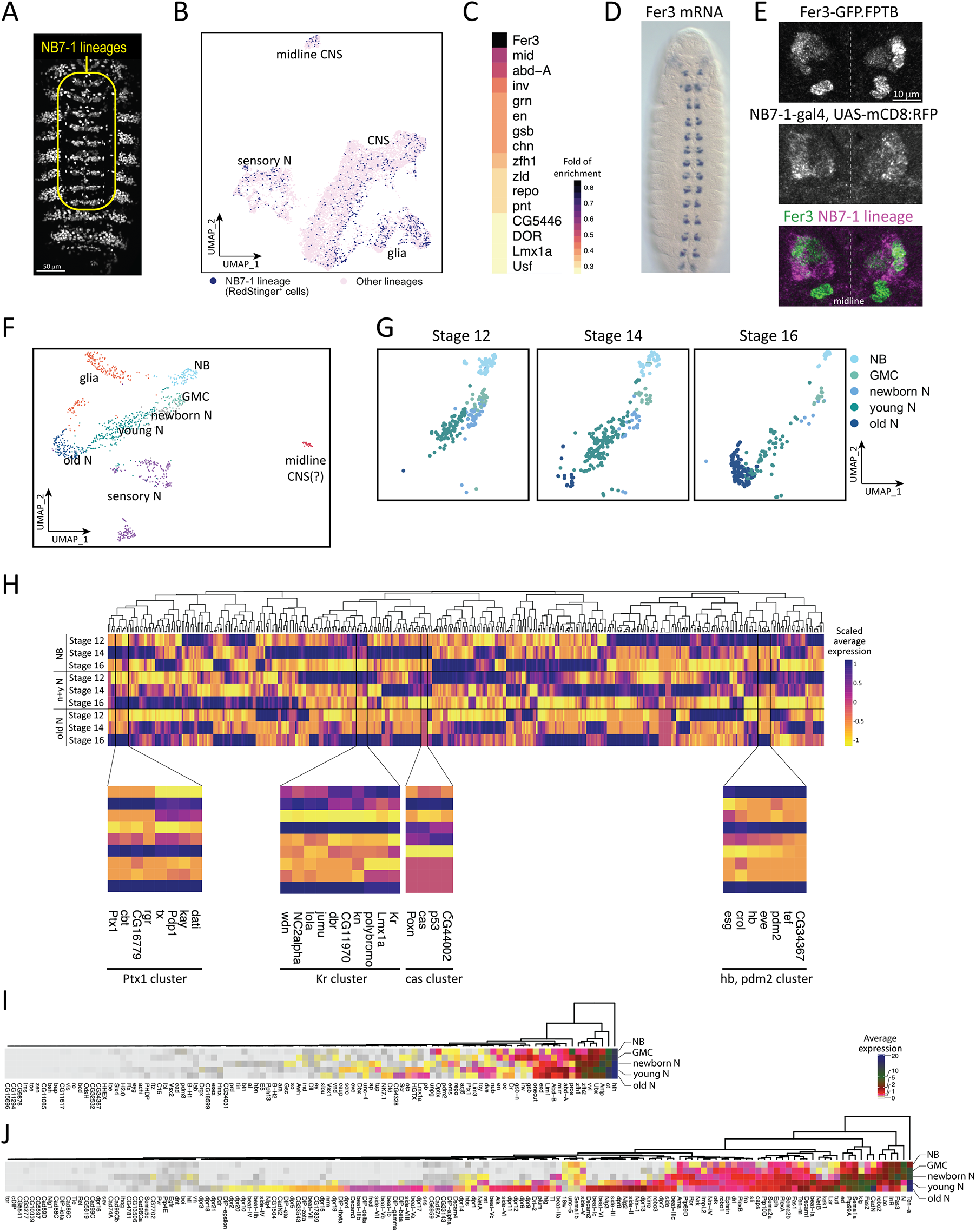
Identification of NB7-1 lineage specific markers and candidate temporal transcription factors. (A) NB7-1-gal4 drives the expression of RedStinger in the whole embryo; yellow outline shows the segmentally repeated NB7-1 lineage, whereas expression outside the outline are epidermal cells that are not part of the neural population. (B) Distribution of NB7-1 lineage (*RedStinger*+) cells in the whole embryo CNS atlas. (C) Heatmap of statistically enriched transcription factors in the NB7-1 lineage. Colors encode the logarithm-transformed folds of enrichment of average expression. (D-E) Expression of Fer 3 mRNA (D) at stage 13 embryo (rotated from https://insitu.fruitfly.org/insitu_image_storage/img_dir_118/insitu118808.jpe) and GFP-tagged Fer3 (E). Fer3 (E, top panel) and NB7-1-gal4 driven expression of membrane-bound RFP (E, middle panel) overlaps in NB7-1 lineage (E, bottom panel). (F) UMAP of reclustered NB7-1 lineage cells. (G) UMAP of NB7-1 neural cells at stage 12, stage 14, and stage 16. NB, neuroblast; GMC, ganglion mother cell; N, neuron. (H) Scaled average expression of transcription factors in neuroblasts (NB), newborn and young neuron (newborn & young N), and mature neuron (old N) at stage 12, stage 14, and stage 16. Color encodes the levels of scaled average expression of each transcription factor at different stages. The bottom panels show the magnified branches of cluster tree clustered with known temporal identify factors (*hb*, *Kr*, *pdm2*, and *cas*), and *Ptx1*, and each cluster is bordered by the vertical black lines. Gene names are listed at the bottom. (I-J) Heatmap of average expression of homeodomain transcription factors (HDTFs) and cell surface molecules (CSMs) in NB7-1 neuroblast (NB), GMC (ganglion mother cells), newborn neurons (newborn N), young neurons (young N) and mature neurons (old N). Gene names are shown at the bottom. Dendrogram at the top shows the clustering of HDTFs (I) and CSMs (J). Colors encode the levels of average expression.

(1) Surprisingly, when unsupervised clustering was performed on all CNS cells,the NB7-1 lineage did not cluster distinctly away from other lineages in UMAP space (Figure 7B). This indicates significant overlap in gene expression patterns between individual NBs.

(2) We were able to identify several genes enriched in NB7-1 (Figure 7C), including some known to be expressed in NB7-1, such as the tandem *engrailed (en)/invected* (inv) genes (Broadus et al., 1995), *gooseberry (gsb)* (Broadus et al., 1995) and *neuromancer 2* (Flybase: *mid)* (Leal et al., 2009). In addition, the top enriched gene was *Fer3*, a transcription factor that has not been previously characterized in the CNS. The *Fer3* RNA expression pattern shows a segmentally repeated cluster of cells adjacent to the midline consistent with expression in NB7-1 (https://insitu.fruitfly.org/cgi-bin/ex/report.pl?ftype=3&ftext=LD04689-a); Figure 7D). We use an endogenous Fer3:GFP transgene and found it overlaps with NB7-1-gal4 UAS-RFP expression, thereby validating its expression in the NB7-1 lineage (Figure 7E).

(2) We then reclustered the 655 NB7-1 lineage cells (*RedStinger*+) from the whole CNS (8595 cells), and found that the NB7-1 lineage generates multiple cell types (Figure 7F). We also found that the NB7-1 lineage shows the same pattern of differentiation seen in the general CNS population (Figure 7G; Table 2).

**Table 2.** Cell types in NB7-1 lineage identified by specific gene expression across development.

|  | Stage 12<br>(187 cells) | Stage 14<br>(229 cells) | Stage 16<br>(205 cells) |
| --- | --- | --- | --- |
| NB | 25.1% | 13.1% | 7.3% |
| GMC | 15.0% | 9.7% | 5.4% |
| Newborn N | 21.4% | 10.0% | 2.9% |
| Young N | 38.0% | 60.3% | 30.7% |
| Old N | 0.5% | 7.0% | 53.7% |

(3) We found genes that cluster with the temporal TFs Hb, Kr, Pdm, or Cas. The genes (*crol*, *eve*, *esg*, and *tef*) are expressed in a similar spatiotemporal pattern to *hb* and *pdm2* (Figure 7H); *dbr, kn,Lmx1A*, and *wdn*, are expressed similarly to *Kr* (Figure 7H), and *CG44002* and *Poxn* similarly to *cas* (Figure 7H). We also found a gene module (Ptx1 cluster, Figure 7H) which shows expression at later timepoints than *hb*, *Kr*, and *cas*; this module may include novel temporal TFs which function late in the lineage. Some of these are targets of temporal TFs in the NB7-1 lineage (e.g. *eve*; (Isshiki et al., 2001)), whereas others may be acting in parallel or after the known neuroblast temporal TFs. In any case, these co-clustered genes are excellent candidates for regulating neuronal identity in the NB7-1 lineage.

(2) We then reclustered the 655 NB7-1 lineage cells (*RedStinger*+) from the whole CNS (8595 cells), and found that the NB7-1 lineage generates multiple cell types (Figure 7F). We also found that the NB7-1 lineage shows the same pattern of differentiation seen in the general CNS population (Figure 7G; Table 2).

### Homeodomain transcription factors and cell surface molecules are upregulated from progenitors to neurons in the NB7-1 lineage

We have shown earlier that homeodomain TFs and CSMs are differentially expressed in post-mitotic motor neurons and interneurons. It remains unclear if these molecules, essential for neuron fate specification and morphogenesis, are inherited from progenitor cells or synthesized de-novo in post-mitotic neurons. Here we use the NB7-1 lineage to address this concept. We computed the average expression of all homeodomain TFs and selected CSMs (see Figure 6) in the NB7-1 lineage. We found that some homeodomain TFs may initially be expressed in NBs; these include spatial factors (*gsb*) or neuroblast self-renewal factors (*pros*). Some homeodomain TFs are modestly expressed in neuroblasts, but upregulated in newborn neurons (*ems*, *pdm2*, and *repo*). Some are highly expressed in young neurons (*acj6, caup, Dbx, eve*, and *scro*), while some homeodomain TFs only expressed at significant levels in mature neurons (*CG4328, Dfd*, and *Nk7.1*) (Figure 7I, Supplemental Figure 1). Most of these CSMs are required for neuron pathfinding and synapse formation, where expression is generally restricted to mature neurons. Interestingly, some CSMs, especially that play a role in axon guidance, show early enrichment in neuroblasts (*Fas3, Sema1a, Ten-a*, and *robo2*) (Figure 7J, Supplemental Figure 1); their function in progenitors remains unknown. In conclusion, we found several homeodomain TFs and CSMs with known roles in neuronal fate determination and morphogenesis to be enriched in the NB7-1 lineage as early as in neuroblasts.

## Methods

### Fly Stocks

Male and female *Drosophila melanogaster* were used. The chromosomes and insertion sites of transgenes (if known) are shown next to genotypes. Previously published gal4 lines, mutants and reporters used were: *NB7-1-gal4^KZ^*(II) (Seroka and Doe, 2019), called *NB7-1-gal4* here; *UAS-RedStinger* [RRID:BDSC_8547]; *UAS-mCD8:RFP*[RRID:BDSC_32218]; and *Fer3-GFP.FPTB*[RRID:BDSC_66447].

### Immunostaining and imaging

DyLight 488-conjugated goat anti-GFP antibody was used (Novus Biologicals, Centennial, CO). Embryos were fixed and stained as previously described (Seroka and Doe, 2019). The samples were mounted in Vectashield (Vector Laboratories, Burlingame, CA). Images were captured with a Zeiss LSM 900 confocal microscope with a *z*-resolution of 0.5 μm. Images were processed using the open-source software FIJI. Figures were assembled in Adobe Illustrator (Adobe, San Jose, CA).

### Embryo dissociation

Cell dissociates were prepared from 8-9 hr (stage 12), 10-11 hr (stage 14) and 15-16 hr (stage 16) embryos respectively. Embryos were washed in DI water, before surface sterilization in 100% bleach for 5 minutes. Embryos were homogenized in Chan-Gehring (C+G) media by six to eight strokes of a loose-fitting dounce. The cell suspension was filtered through a 40 μm Nitex mesh, and cells were pelleted in a clinical centrifuge at 4°C (setting 5, IEC). The cell pellet was washed twice by pouring off the supernatant and gently triturating the pellet in fresh C+G. Percent cell-survival was determined for each dissociate by BioRad TC-20 trypan-blue assay.

### Generating raw data

Sample preparation was performed by the University of Oregon Genomics and Cell Characterization core facility (https://gc3f.uoregon.edu/) Dissociated cells were run on a 10X Chromium platform using 10X V2 chemistry targeting 10,000 cells per sample. Following cDNA library preparation, the library for each timepoint was amplified with 15 cycles of PCR before sequencing on two separate Illumina Hi-seq lanes, providing two technical replicates for each timepoint (stages 12, 14, 16). Replicates were merged using the CellRanger Aggregate function prior to quality control and downstream analysis. Reads were aligned to the Drosophila genome (BDGP6.22) and protein coding reads were counted. The resulting sequencing data were analyzed using the 10X CellRanger pipeline, version 3.1.0 (Zheng et al., 2017) and the Seurat software package for R, v3.1.2 using standard quality control, normalization, and analysis steps. Cells were filtered by % expression of mitochondrial genes, indicating high stress state. Only cells expressing <20% mitochondrial reads were retained for analysis. Additionally, cells containing reads for <50 and >3000 unique genes were filtered out of downstream analysis. For each gene,expression levels were normalized by total expression, multiplied by a scale factor (10,000) and log-transformed. Differential expression analysis was performed with the FindAllMarkers function in Seurat using Wilcoxon rank sum test. Tissue identity was determined by the expression score of tissue-specific genes (Supplemental Table 2) with the AddModuleScore function in Seurat. Cells were subsetted for further analysis based on the clustering and expression of ground-truth genes (see Results sections).

## Abbreviations

(sc-RNAseq): Single cell RNA sequencing
(Eve): Even-skipped
(TTF): Temporal transcription factor
(Hb): Hunchback
(Kr): Kruppel
(Pdm): Nubbin/Pou domain 2
(Cas): Castor
(NB7-1-gal4): NB7-1 split gal4
(CNS): central nervous system.

## Ethical Approval and Consent to participate

Not applicable; no vertebrate or human subjects.

## Consent for publication

All authors have approved this manuscript.

## Availability of data and materials

All code used for analysis will be uploaded to GitHub upon acceptance. No new fly stocks were generated, and all fly stocks are available from public stock centers or by request.

## Author contributions

A.Q.S., S.-L.L and C.Q.D. conceptualized the work. A.Q.S. and S.-L.L. performed experiments and analyzed results. All authors contributed to writing the manuscript.

## Acknowledgements

We thank Noah Dillon and Dylan Farnsworth for comments on the manuscript. We thank Maggie Weitzman in the Genomics Core for library preparation and sequencing.

## Authors information

Howard Hughes Medical Institute, Institute of Neuroscience, University of Oregon, Eugene 97403, USA.

---

**Supplemental Table 1. Genes enriched in each embryonic tissue cluster in Figure 1.**

A spreadsheet of markers of each individual cluster determined by FindAllMarkers function in Seurat. Identity of each cluster is determined by tissue-specific genes (Supplemental Table 2).

**Supplemental Table 2. Genes used as tissue-specific “ground truth” in Figure 1.**

Sixteen separate sheets are included. Each sheet contains a list of genes that are annotated to be expressed in the tissue based on in situ hybridization database (https://insitu.fruitfly.org/cgi-bin/ex/insitu.pl), and used to identify tissue-specific clusters in Figure 1.

**Supplemental Table 3. Genes enriched in each CNS tissue cluster.**

A spreadsheet of markers of each individual cluster determined by FindAllMarkers function in Seurat. Identity of each cluster is determined by ground-truth genes; see Figure 4.

**Supplemental Figure 1.**
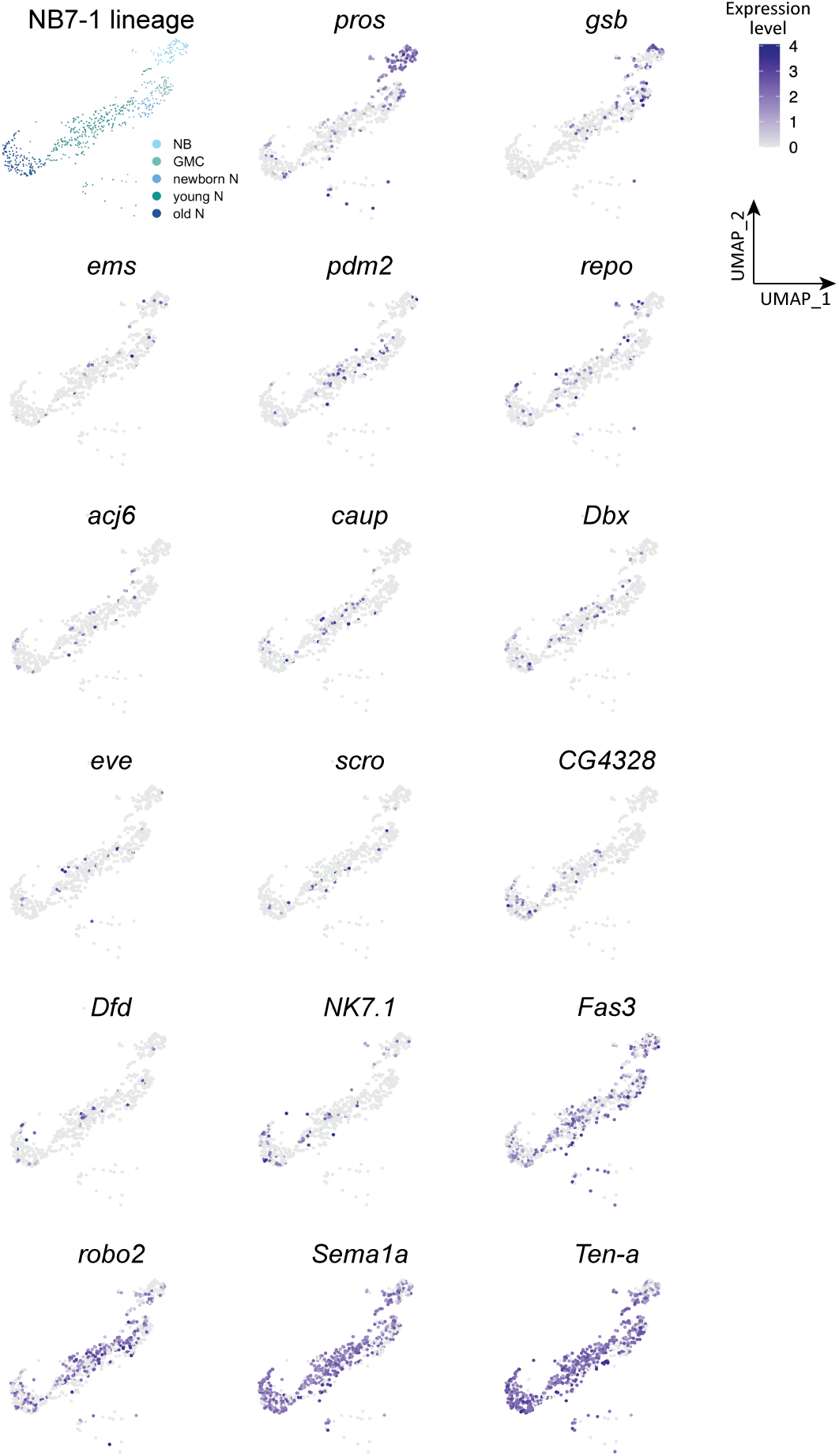
Genes expressed in NB7-1 lineage. Atlas of NB7-1 lineage in UMAP and feature plots of genes expressed in NB7-1 lineage. Colors encode the expression level.

